# Widespread latitudinal asymmetry in marginal population performance

**DOI:** 10.1101/529560

**Authors:** Fernando Pulido, Bastien Castagneyrol, Francisco Rodríguez-Sánchez, Yónatan Cáceres, Adhara Pardo, Eva Moracho, Johannes Kollmann, Fernando Valladares, Johan Ehrlén, Alistair S. Jump, Jens-Christian Svenning, Arndt Hampe

**Affiliations:** Institute for Dehesa Research (INDEHESA), University of Extremadura, Plasencia, Spain; BIOGECO, INRA, Univ. Bordeaux, 33610 Cestas, France; Estación Biológica de Doñana (EBD-CSIC), Sevilla, Spain; Departamento de Biología Vegetal y Ecología, Universidad de Sevilla, Avda. Reina Mercedes s/n, 41012 Sevilla, Spain; School of Life Sciences, Technical University of Munich, 85350 Freising, Germany; Museo Nacional de Ciencias Naturales (MNCN-CSIC), Madrid, Spain; Department of Ecology, Environment and Plant Sciences, and Bolin Centre for Climate Research, Stockholm University, 10691 Stockholm, Sweden; Biological & Environmental Sciences, Faculty of Natural Sciences, University of Stirling. Stirling, FK9 4LA, UK; Center for Biodiversity Dynamics in a Changing World (BIOCHANGE) and Section for Ecoinformatics and Biodiversity, Department of Biology, Aarhus University, Ny Munkegade 114, 8000 Aarhus C, Denmark

**Keywords:** centre-periphery hypothesis, demographic rates, latitudinal asymmetry, marginal populations, population performance, range margin, range shift

## Abstract

**Aim:** Range shifts are expected to occur when populations at one range margin perform better than those at the other margin, yet no global trend in population performances at range margins has been demonstrated empirically across a wide range of taxa and biomes. Here we test the prediction that, if impacts of ongoing climate change on population performance are widespread, then populations from the high-latitude margin (HLM) should perform as well as or better than central populations, whereas populations at low-latitude margins (LLM) populations should perform worse.

**Location:** Global

**Time period:** 1898–2020

**Major taxa studied:** Plants and animals

**Methods:** To test our prediction, we used a meta-analysis quantifying the empirical support for asymmetry in the performance of high- and low-latitude margin populations compared to central populations. Performance estimates were derived from 51 papers involving 113 margin-centre comparisons from 54 species and 705 populations. We then related these performance differences to climatic differences among populations. We also tested whether patterns are consistent across taxonomic kingdoms (plants *vs*. animals) and across habitats (marine *vs*. terrestrial).

**Results:** Populations at margins performed significantly worse than central populations and this trend was primarily driven by the low-latitude margin. Although the difference was of small magnitude, it was largely consistent across biological kingdoms and habitats. The differences in performance were positively related to the difference in average temperatures between populations during the period 1985–2016.

**Major conclusions:** The observed asymmetry in marginal population performance confirms predictions about the effects of global climate change. It indicates that changes in demographic rates in marginal populations, despite extensive short-term variation, can serve as early-warning signals of impending range shifts.

## Introduction

Ongoing climate changes are predicted to increase mismatches between current environmental conditions and the climate to which local populations are adapted (Svenning & Sandel 2013). These mismatches should in turn result in range-wide asymmetries in population growth rates with positive rates at the upper latitudinal or altitudinal range edges, and negative ones at low-latitude or altitude edges. Such asymmetries in population growth rates have been hypothesized to be the main driver of large-scale geographical range shifts (Parmesan et al. 1999, Sexton et al. 2009, Lenoir & Svenning 2015). Yet, we know little about how generalised asymmetries in marginal population growth rates are globally. Population growth rates are difficult to monitor directly, but the demographic processes underlying these rates, such as survival and fecundity, are more accessible to short-term observation. Quantifying the global extent of asymmetry in demographic processes should thus allow us to assess existing disequilibrium of species ranges with climate and hence the propensity of species to shift their range. Such knowledge is crucial to accurately forecast future climate-driven range shifts (Dullinger et al. 2012, Normand et al. 2013) and changes in ecosystem functioning, and for informing resource and conservation planning.

Putatively climate-driven distribution changes have been extensively reported in recent years (Chen et al. 2011, Wiens 2016, Lenoir et al. 2020). However, causal relationships with recent climate changes are difficult to establish because range limits can also be constrained by diverse non-climatic factors, such as habitat availability, dispersal limitation or biotic interactions (Louthan et al. 2015, Hargreaves et al. 2014, Lee-Yaw et al. 2016, Pironon et al. 2017), and especially long-lived and immobile species may accumulate extensive extinction debts and colonisation credits (Talluto et al. 2017). Changes in the performance of marginal populations should hence represent a much more direct and immediate indicator of species’ response to climate warming than distribution changes (Vilà-Cabrera et al. 2019). Still, the effects of climate on population performance will often be difficult to detect except in meteorologically extreme years. Long-term observations that enable detecting such events in marginal population dynamics are, however, rare, especially for populations at contracting range margins (Hill et al. 2011, James et al. 2015, Fredston-Hermann et al. 2020, Hastings et al. 2020, Kunstler et al. 2021). Hence, we need other approaches to assess if range-wide asymmetries in population performances occur.

Here, we use the abundant empirical literature spawned by the so-called centre-periphery (CP) paradigm to examine differences in population performance between the range centre and the high- and low-latitude margins for a wide range of taxa. This paradigm states that the size, density and long-term growth rate of populations tend to decrease from the centre towards the periphery of the range as environmental conditions become increasingly less favourable (Brown 1984, Sagarin & Gaines 2006, Sexton et al. 2009) (Fig. 1). The centre-periphery paradigm has motivated hundreds of comparisons of various indicators of population performance (including measures of individual survival or fecundity, population viability and others) in central and marginal populations (Pironon et al. 2017).

**Figure 1.**
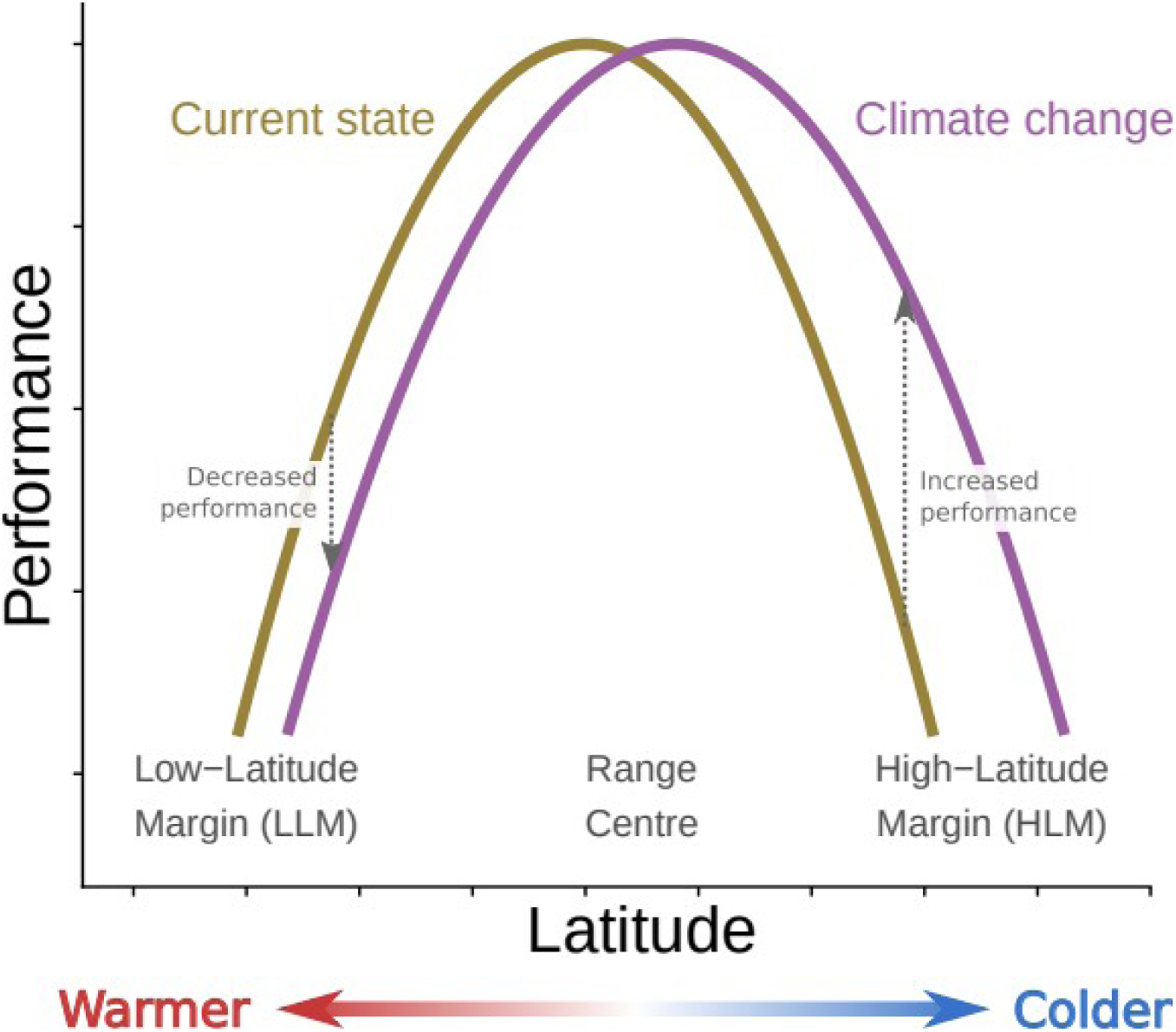
The centre-periphery hypothesis postulates that population performance is maximal around the range centre and decreases towards the margins of the distribution range, as environments become less suitable. Under current climate change, the optimal climate zones would displace polewards so that high-latitude populations (HLM) would increase their fitness whereas low-latitude populations (LLM) experience a decrease. Hence, the difference in performance between high-latitude and central populations would reduce with climate change, while low-latitude populations would show greater differences to central populations.

We selected a comprehensive sample of published studies to compare measures of population performance in sites located at the centre and at the high-latitude margins (HLM) or low-latitude margins (LLM) of species ranges. We predicted that if impacts of ongoing climate change on population performance are widespread, then (i) HLM populations should perform as well as or better than central populations, whereas LLM populations should perform worse than central and HLM populations (Fig. 1). To test this prediction, we employed information from empirical studies to quantify differences in the performance of HLM, LLM and central populations, and to test if patterns are consistent across taxonomic kingdoms (plants *vs*. animals) and across habitats (marine *vs*. terrestrial). We also predicted that if climate is a major driver of differences in population performance, then (ii) performance differences should increase with the difference in climate between central and marginal populations (Fig. 1). To test this prediction, we relate the observed differences in performance between central and peripheral populations to differences in climate.

## Materials and methods

### Data compilation

We searched Web of Knowledge^®^ and Scopus until 19^th^ April 2020 for publications in peer-reviewed international scientific journals using key search terms in the title or the abstract. In addition, we searched Google Scholar using the same terms in the full text of scientific publications, and restricting our selection to the first 200 references found. The terms ‘centre/center-periphery’, ‘central-marginal’, ‘abundant centre/center’, and ‘latitudinal cline’ were introduced in combination with performance related terms including ‘fecundity’, ‘performance’, ‘survival’, ‘recruitment’ and ‘population growth rate’. Three filters were then applied to the initial subset of papers. First, we only considered studies reporting field data from natural populations, including control populations of transplant experiments if these were measured at their home sites and met all other criteria. Second, we only considered studies with at least two central and two peripheral populations (i.e., true replicates). Third, we only considered papers that provided sufficiently clear criteria for the definition of central and peripheral range parts relative to the global range of the target species (and not only parts of it). This filtering procedure resulted in a total of 51 publications, with 113 centre-periphery contrasts of 54 species. The workflow and output of our compilation and selection process is described in Supplementary Information 1.

We extracted the reported performance metrics from each primary paper and assigned them to one of three categories: (i) ‘Survival’ (e.g., mortality of individuals or ramets, rates of fruit abortion or germination), (ii) ‘Reproduction’ (e.g., proportion of actively reproducing individuals, seed number, gonadal mass, total seed or egg mass), or (iii) ‘Lifetime fitness’ (e.g., different estimates of population growth rate). Moreover, we assigned each case study to one of two major taxonomic groups (plants *vs*. animals) and habitat (terrestrial *vs*. marine). Two major kinds of papers provided suitable information: (i) explicit CP comparisons of mean performance values from populations classified as central or marginal by the authors, and (ii) papers reporting on latitudinal clines. In the first case, we followed the criteria of the original authors for classifying populations as central or marginal. In the second case, we selected the three most central and the three most marginal populations along the gradient (rarely more if several populations were located closely together). We extracted quantitative data for our target metrics either manually from text and tables or from figures with Dagra digitizing software version 2.0.12 (Blue Leaf Software 2016). We recorded mean values for each individual population (Fig. S3), and then calculated the average performance, sample size and resulting standard deviation for central, high- and low-latitude margins, respectively.

### Meta-analysis of differences between marginal and central populations

#### Effect Sizes

We used Hedges’ *d* statistic as our standardised measure of effect size. Hedges’ *d* is the most appropriate effect size to compare raw means when both positive and negative values are present in data (Koricheva et al. 2013). Hedges’ *d* was calculated as:

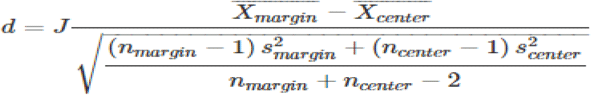

where

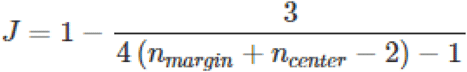

and 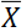, n and s^2^ the mean, sample size and sampling variance.

Negative values of *d* indicate lower performance in marginal (either HLM or LLM) populations than in central populations (consistent with the CP paradigm), whereas positive values indicate higher performance. The sampling variance of effect sizes was:

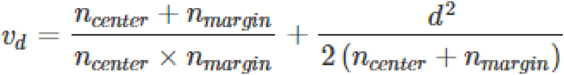

Note that *v*_*d*_ contains information about both the sample size and the standard deviation (within *d*^*2*^) of the original studies; it hence can be used to weight the relative importance of studies within the meta-analysis. In some papers, both HLM populations and LLM populations were compared to the same central populations, resulting in an overestimated pooled sample size (*N = n*_*center*_ + *n*_*margin*_) because, for such primary papers, *n*_*center*_ is counted twice. We manually corrected *N* in all such cases before conducting the analysis.

#### Meta-analytical models

Our dataset had a hierarchical structure as some primary papers contained several case studies. We accounted for this potential non-independence of cases by estimating model heterogeneity from multiple sources: (i) among true effect sizes, (ii) among CP comparisons stemming from the same primary papers (by computing the variance-covariance matrix among all effect sizes), and (iii) among groups of moderators. This was done using multi-level error meta-analysis with the *rma*.*mv* function of the R package *metafor* v. 2.0-0 (Viechtbauer 2010). Primary paper identity was declared as a random factor and individual CP comparisons were nested as random factor within primary papers. We estimated variance components for primary papers (σ_1_^2^) and case studies (σ_2_^2^) together with intra-class correlations (ρ), i.e., correlations between true effect sizes from the same study (such that ρ= σ_1_^2^ + σ_2_^2^).

We first calculated grand mean effect size as the overall weighted mean across all effect sizes (Borenstein et al. 2007). This corresponds to a random-effects meta-analysis, where heterogeneity among true effect sizes (*τ*^2^) is used to weight individual effects sizes (weight = 1/(*v* + τ^2^)). Then, we used multi-level (hierarchical) meta-analyses to test the effect of three moderators: Margin (HLM vs. LLM), Kingdom (animals vs. plants) and Habitat (marine vs. terrestrial). We built a set of the 17 possible models including all possible combinations of simple effects (*n* = 7 models) and two-way interactions among Margin, Kingdom and Habitat (*n* = 10 models). We ranked these 17 models plus the null model (i.e., intercept only) according to their AIC_c_ using the R package *glmulti* v. 1.0.7 (Calcagno 2013). For each model, we calculated ΔAIC_c_ and AIC_c_ weight (*w*_*i*_). Models within ΔAIC_c_ < 2 were considered as best models, given the data structure and the model set (Table 1). AIC_c_ weights represent the probability that a given model is the best model in the set of models considered. For each moderator, we then estimated its relative importance (*w*_*H*_) by summing all *w*_*i*_ of the models including this moderator (*w*_*H*_ = Σ*w*_*i*_); *w*_*H*_ can be interpreted as the probability that a given moderator is included in the best model (Fig. S2). Finally, we estimated model parameters for all competing models with ΔAICc < 2. We reported model parameter estimates for the best model and, whenever necessary, for competing models. Further details upon the meta-analysis, including several assessments of its inherent reliability (e.g., publication bias, balanced representation of moderators, etc.) are shown in Supplementary Information 2.

**Table 1.**
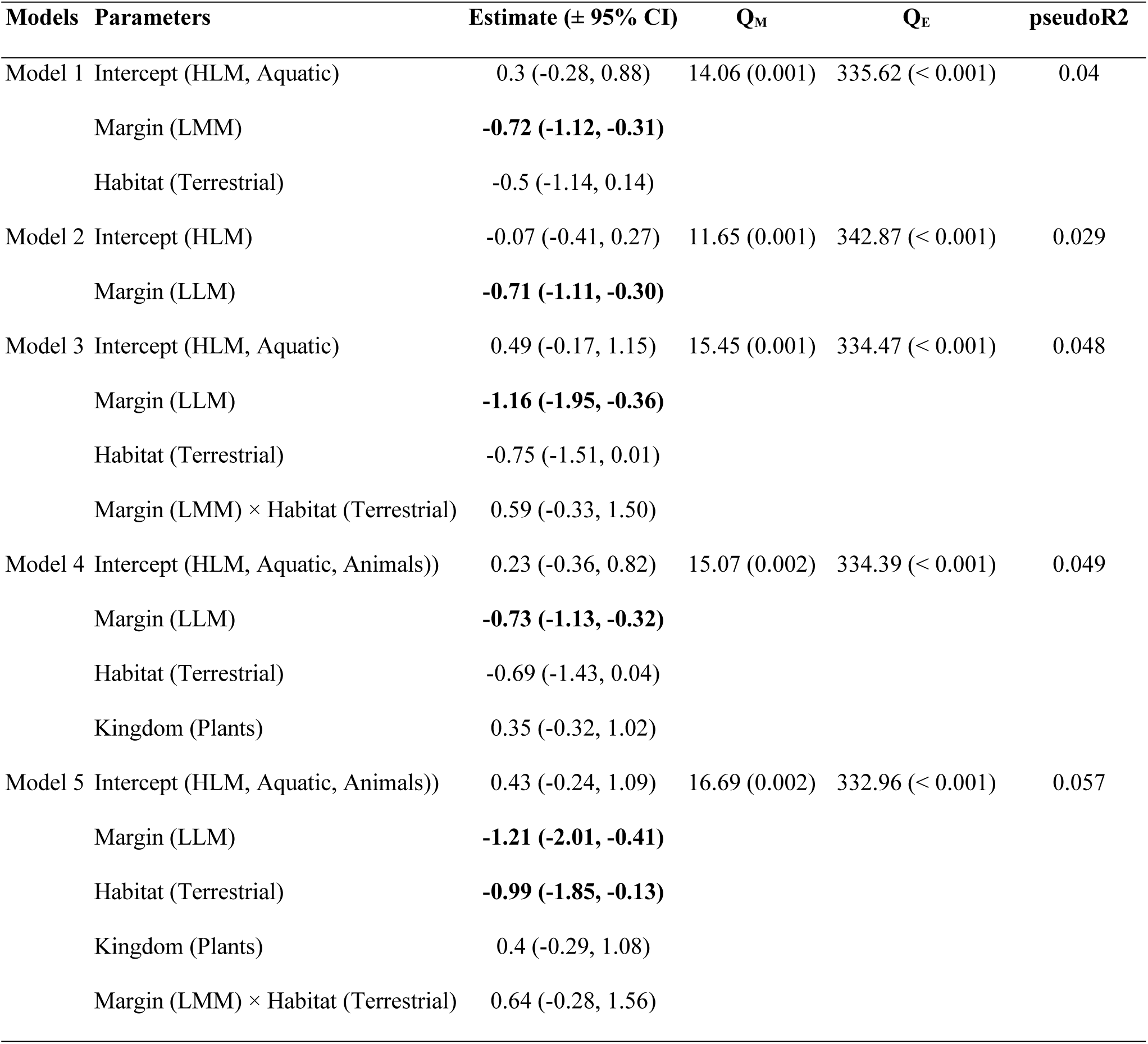
Summary of the five models retained in the set of best models (i.e., with ΔAIC_c_ < 2). Margin explained a significant amount of heterogeneity in each of the five competing best models whereas neither Kingdom nor Habitat explained a significant amount of heterogeneity in any of the five models retained in the set of best models. QM and associated P-values represent the test associated with each moderator, separately. Pseudo R^2^ were calculated as 1 – LLR, where LLR is the ratio between the log-likelihood of model i and the log-likelihood of the null model.

| Models | Parameters | Estimate (± 95% CI) | Q<sub>M</sub> | Q<sub>E</sub> | pseudoR<sup>2</sup> |
| --- | --- | --- | --- | --- | --- |
| Model 1 | Intercept (HLM, Aquatic) | 0.3 (-0.28, 0.88) | 14.06 (0.001) | 335.62 (< 0.001) | 0.04 |
|  | Margin (LMM) | **-0.72 (-1.12, -0.31)** |  |  |  |
|  | Habitat (Terrestrial) | -0.5 (-1.14, 0.14) |  |  |  |
| Model 2 | Intercept (HLM) | -0.07 (-0.41, 0.27) | 11.65 (0.001) | 342.87 (< 0.001) | 0.029 |
|  | Margin (LLM) | **-0.71 (-1.11, -0.30)** |  |  |  |
| Model 3 | Intercept (HLM, Aquatic) | 0.49 (-0.17, 1.15) | 15.45 (0.001) | 334.47 (< 0.001) | 0.048 |
|  | Margin (LLM) | **-1.16 (-1.95, -0.36)** |  |  |  |
|  | Habitat (Terrestrial) | -0.75 (-1.51, 0.01) |  |  |  |
| | Margin (LMM) × Habitat (Terrestrial) | 0.59 (-0.33, 1.50) | | | |
| Model 4 | Intercept (HLM, Aquatic, Animals)) | 0.23 (-0.36, 0.82) | 15.07 (0.002) | 334.39 (< 0.001) | 0.049 |
|  | Margin (LLM) | **-0.73 (-1.13, -0.32)** |  |  |  |
|  | Habitat (Terrestrial) | -0.69 (-1.43, 0.04) |  |  |  |
|  | Kingdom (Plants) | 0.35 (-0.32, 1.02) |  |  |  |
| Model 5 | Intercept (HLM, Aquatic, Animals)) | 0.43 (-0.24, 1.09) | 16.69 (0.002) | 332.96 (< 0.001) | 0.057 |
|  | Margin (LLM) | **-1.21 (-2.01, -0.41)** |  |  |  |
|  | Habitat (Terrestrial) | **-0.99 (-1.85, -0.13)** |  |  |  |
|  | Kingdom (Plants) | 0.4 (-0.29, 1.08) |  |  |  |
| | Margin (LMM) × Habitat (Terrestrial) | 0.64 (-0.28, 1.56) | | | |

### Relationship between climate and differences in population performance

#### Climate data

We gathered the geographical coordinates of all populations included in the meta-analysis from the primary papers (*n* = 705 populations). For each population, we calculated the average annual temperature between 1985 and 2016 (i.e., when most studies were performed) based on monthly temperature data from CRU TS 4.04 (Harris et al. 2020) for terrestrial species and HadISST 1.1 (Rayner et al. 2003) for marine species. For terrestrial taxa, we also extracted average annual precipitation at each site, again from CRU TS 4.04. We could not match climate data for two fish species (Heibo et al. 2005, Power et al. 2005; Appendix 1) and hence excluded these species from the analyses. The final dataset for the climatic analysis contained 683 populations from 52 species (37 terrestrial, 15 marine) and 109 margin–centre comparisons (Fig. S3). We then aggregated populations to calculate average temperature and precipitation for each combination of study, species, performance variable, and region (either central, HLM, or LLM). We could then relate each comparison of performance between a margin (HLM or LLM) and the central range (i.e., Hedges’ *d*) with the difference in average temperatures or precipitation between both regions.

#### Analysis of relationships between climate and population performance

To assess the relationship between the differences in performance and the differences in climate between marginal and central populations, we used generalised additive mixed models (function gam in the R package *mgcv*, version 1.8-17 [Wood 2006]) using the temperature differences as predictor, and the study as random effect (to control for lack of independence). We weighted performance effect sizes by their variances so that their influence in model calibration was inversely related to their uncertainty. For the terrestrial taxa, we also fitted a similar model including precipitation and its interaction with temperature as predictors (see Supplementary Information 3 for further details).

## Results

Marginal populations performed on average worse than central populations, since grand mean effect size was negative (−0.35; 95% CI: -0.66, -0.04). There was a significant amount of heterogeneity, and 58% of the total heterogeneity was due to among-study heterogeneity (*τ*^2^ = 1.29, *Q*_*E*_ = 362.48, *P <* 0.0001). Five models received relatively strong support, at the level of ΔAIC_c_ ≤ 2.0. All five included margin type as a moderator (*w*_H_ = 0.99).

Performance declined more strongly towards the low-latitude margin (effect size: - 0.94; 95% CI: -1.42, -0.46; estimated from the model with Margin as the sole moderator) than towards the high-latitude margin (effect size: -0.37; 95% CI: -0.80, 0.07) (Fig. 2). The difference between population performance in both margins was significant in the five models with ΔAICc ≤ 2 (Table 1). The best model explained only 5% of the total heterogeneity. The five models with ΔAICc ≤ 2 also included Habitat, Kingdom and the Margin × Habitat interaction as moderators, but their relative importance was low (*w*_H_ < 0.75); only one of the differences between aquatic and terrestrial habitats was significant, whereas none of the comparisons between plants and animals were significant (Fig. 3).

**Figure 2.**
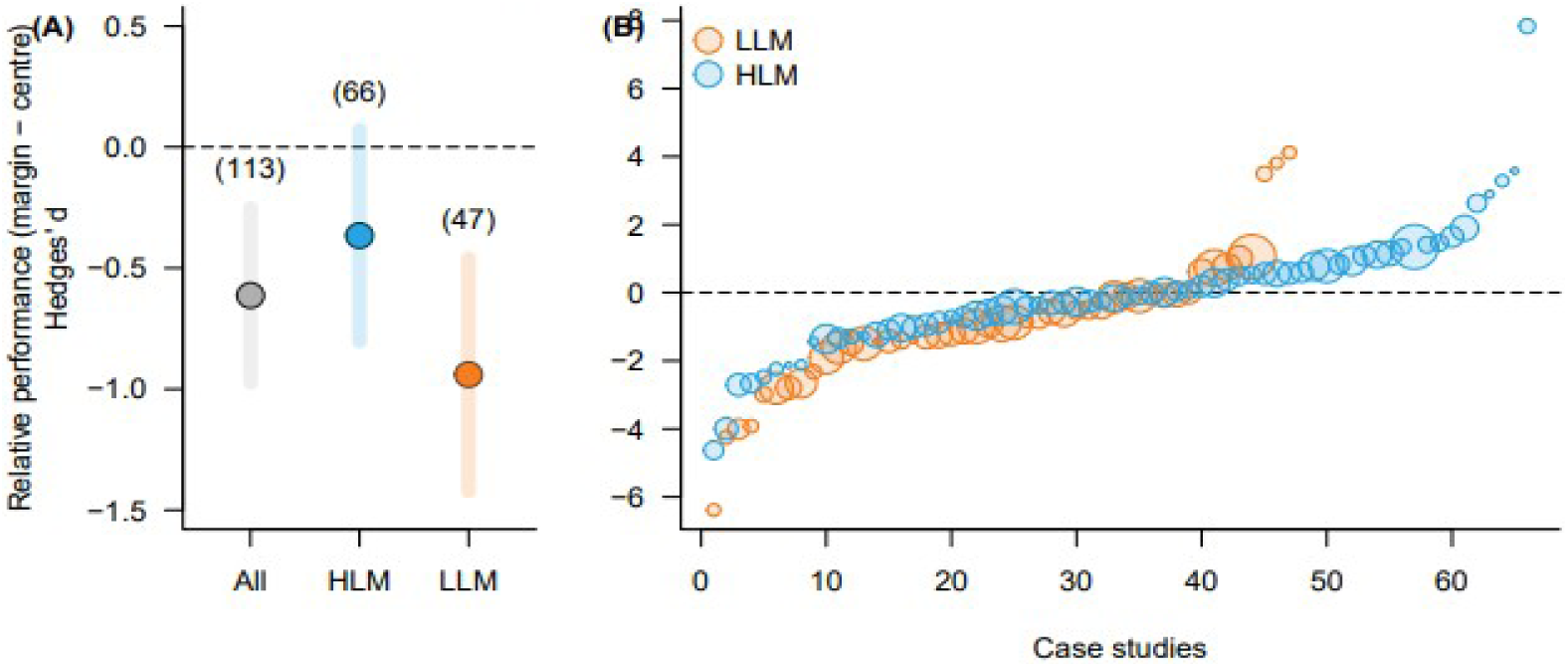
Observed differences in performance (Hedges’*d* effect sizes) between marginal (high-latitude, HLM, and low-latitude, LLM) and central populations across all species and studies. **(A)** Grand mean (grey) and margin-specific (blue and orange) combined effect sizes. Error bars represent 95% confidence intervals. Numbers in parentheses correspond to the number of case studies. **(B)** Individual effect sizes for HLM and LLM case studies ranked from the lowest to the highest value. Dot size is proportional to the weight of individual effect sizes in the meta-analysis. Both in **(A)** and **(B)**, positive and negative values indicate higher and lower performances in marginal than in central populations, respectively. Horizontal dashed lines represent the null hypothesis of no difference in the performance of central vs. marginal populations.

**Figure 3.**
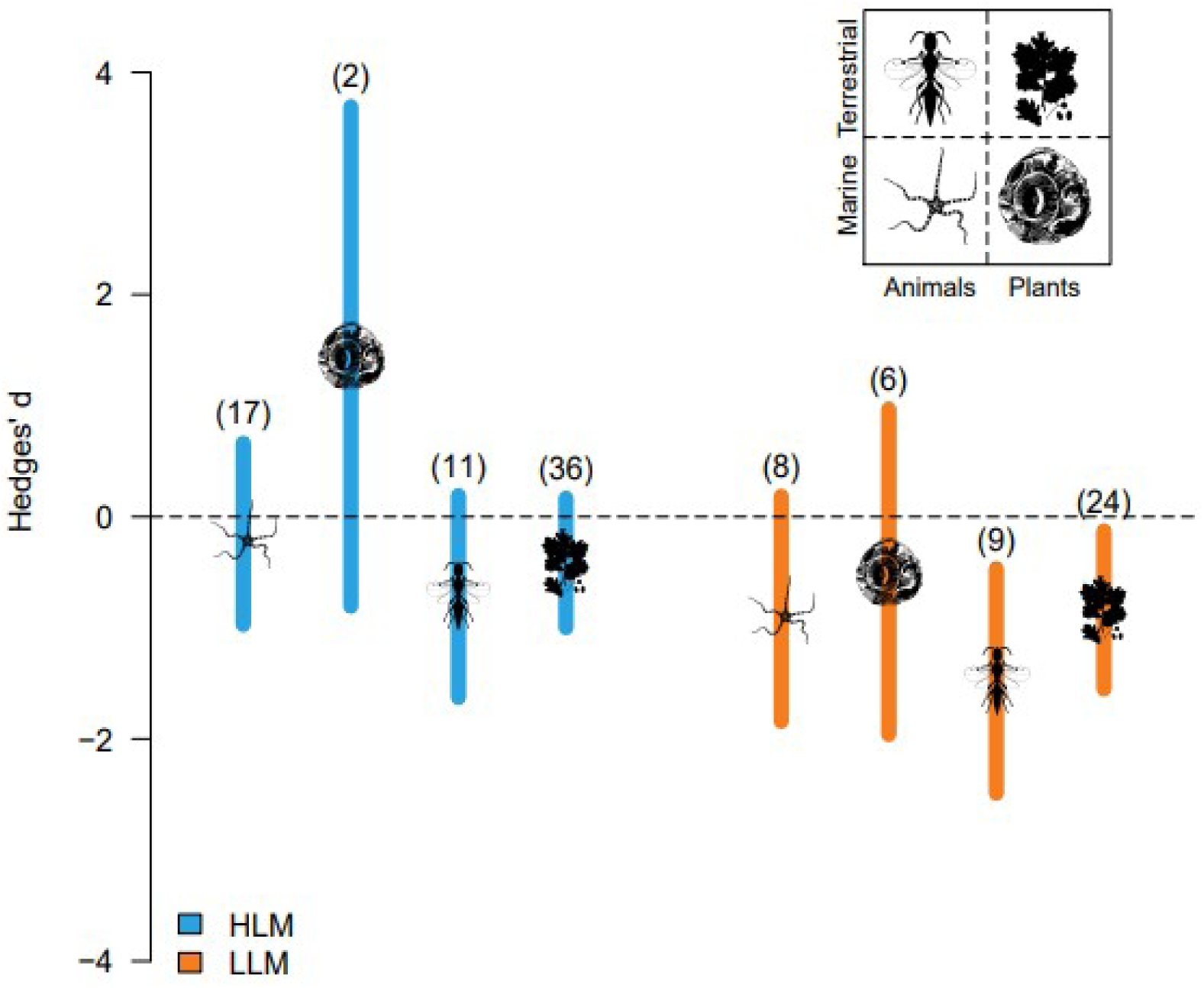
Asymmetry in population performance at High Latitude Margins (HLM) and Low Latitude Margins (LLM) for each Kingdom and Habitat. Symbols representing Habitats and Kingdoms are centred on the mean estimate. Vertical bars represent 95% CI estimated from the multi-level meta-analysis. Negative and positive values indicate lower and higher performance of marginal populations as compared to central populations, respectively. Numbers within parentheses indicate the number of case studies for each category.

The differences in performance between marginal and central populations were significantly related to the difference in their average temperature in the period 1985–2016 (*F* = 3.39, *P* = 0.017; Table S3; total deviance explained by an additive mixed model: 20.8%). As predicted, performance decreased with increasingly departing temperatures from central populations, although the decline was asymmetric between high- and low-latitude populations (Fig. 4): HLM populations experiencing 5 ºC colder temperatures than central populations showed similar performance, whereas LLM populations experiencing 5°C warmer temperatures performed significantly worse (Fig. 4). Differences in population performance were not related to geographical distance between marginal and central populations (Fig. S4). In terrestrial species, population performance showed a similar asymmetric response to temperature (i.e., higher overall performance in HLM than LLM for similar temperature deviations), although there was a significant interactive effect of precipitation (*F* = 2.70, *P* = 0.024, total deviance explained = 29.5%; Table S4). With decreasing precipitation, relative performance in marginal populations decreased faster at both margins, albeit more so in low-latitude populations (Fig. 5).

**Figure 4.** Relationship between the observed difference in performance (Hedges’ *d*) and the difference in average temperatures between peripheral and central populations for the period 1985-2016 for all studied species (52 species: 37 terrestrial and 15 marine; n = 109 margin-centre comparisons). Positive values of Hedges’ *d* indicate better performance in the margin compared to central populations, and vice versa. Point size is inversely related to Hedges’ *d* variance for each contrast (i.e., bigger points represent stronger effect sizes). The curve represents the fit of a generalized additive mixed model (GAMM) with temperature as predictor (and study as random effect to control for lack of independence). The shaded area around the GAMM curve represents the standard error of the prediction. HLM = high-latitude margin, LLM = low-latitude margin.

**Figure 5.**
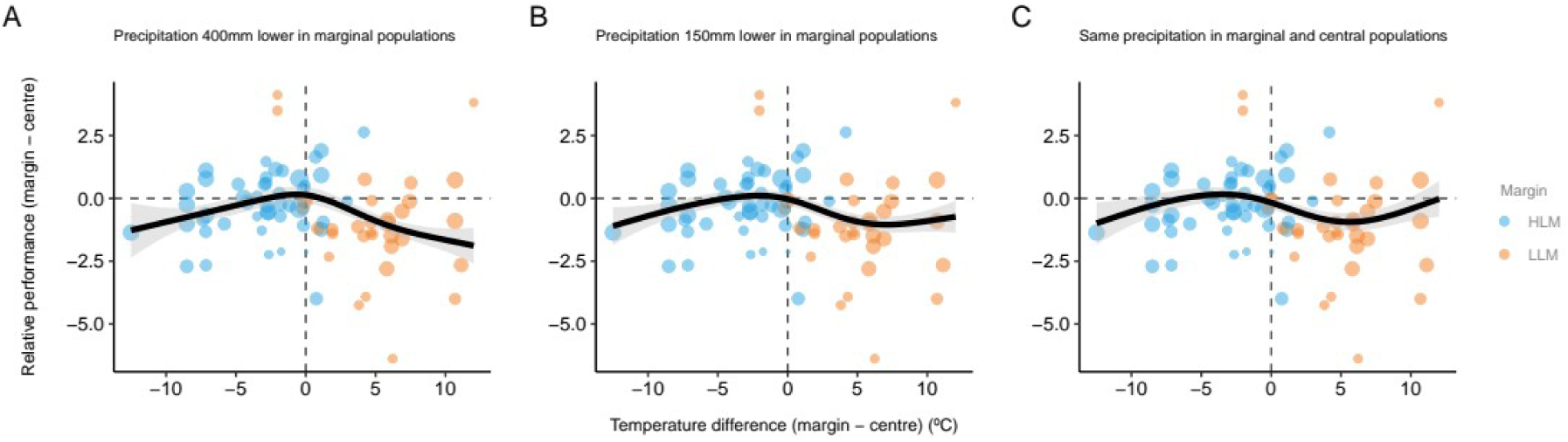
Relationship between the observed difference in performance (Hedges’ *d*) and the difference in average temperatures between peripheral and central populations for the period 1985–2016 for 37 terrestrial species (n = 80 margin-centre comparisons). Positive values of Hedges’ *d* indicate better performance in the margin compared to central populations, and vice versa. Point size is inversely related to Hedges’ *d* variance for each contrast (i.e., bigger points represent stronger effect sizes). The black curve represents the fit of a generalized additive mixed model (GAMM) with temperature, precipitation and their interaction as predictors (and study as random effect to control for lack of independence). The shaded area around the GAMM curve represents the standard error of the prediction. As the interaction between temperature and precipitation was significant, results are shown for three scenarios (annual precipitation 400 mm lower, 150 mm lower, or same in marginal as in central populations). These values approximate the first, second, and third quartiles, respectively, of precipitation differences between marginal and central populations observed in our dataset. HLM = high-latitude margin, LLM = low-latitude margin.

## Discussion

Overall, our results show that populations from the centre of the range tend to outperform those residing at the range margins, and that these differences are more pronounced at low-latitude margins. Such latitudinal asymmetry is predicted when the environmental conditions relevant for population performance are directionally displaced (Lenoir & Svenning 2015; Fig. 1). Global warming has provoked a rapid large-scale poleward displacement of climatic zones since the 1970s, and the trend is predicted to further accelerate through the coming decades (IPCC 2013). The observed difference is therefore likely to largely result from ongoing climate change, although we cannot exclude effects of changes in factors unrelated with current climate, such as historical colonisation lags or biotic interactions (Normand et al. 2011, Hargreaves et al. 2014, Louthan et al. 2015). To achieve further insights into this question, we thoroughly searched the literature (including the BIOSHIFT database, Lenoir et al. 2020) for reports on observed range shifts of the species included in our meta-analysis. We detected a total of 26 range shifts concerning 16 of our species; most shifts (19 out of 26) were polewards (see Table 2). Although limited, this evidence suggests that demographic rates in marginal populations act indeed as early-warning signals of impending range shifts.

**Table 2.** List of species included in the meta-analysis for which information about ongoing range shifts has been reported in the scientific literature. All cases stem from the northern hemisphere. See Appendix 2 for full references.

| Organism | Species | Type of range shift | Reference |
| --- | --- | --- | --- |
| Bird | *Cyanistes caeruleus* | expansion northwards | Massimino et al. 2015 |
|  |  | expansion northwards | Bromer 2004 |
|  |  | shift southwards | Tayleur et al. 2015 |
|  |  | shift northwards | Potvin et al. 2017 |
|  |  | shift northwards | Virkkala & Lehikoinen 2017 |
|  | *Ficedula hypoleuca* | expansion northwards | Virkkala & Lehikoinen 2017 |
|  | *Dendroica caerulescens* | expansion northwards | Zuckerberg et al. 2009 |
| Fish | *Girella elevata* | expansion northwards | Last et al. 2015 |
|  |  | expansion northwards | Stuart-Smith et al. 2010 |
| Herb | *Plantago coronopus* | expansion northwards | Groom 2013 |
| Herb | *Silene acaulis* | expansion south | Groom 2013 |
| Herb | *Cirsium heterophyllum* | expansion northwards | Groom 2013 |
|  | *Cirsium acaule* | expansion northwards | Amano et al. 2014 |
| Herb | *Himantoglossum hircinum* | expansion northwards | Van der Meer et al. 2016 |
|  |  | contraction south | Pfeifer et al. 2010 |
| Herb | *Cypripedium calceolus* | contraction south | Lesica & Crone 2017 |
| Seaweed | *Fucus guiryi* | contraction south | Riera et al. 2015 |
|  |  | contraction south | Lourenço et al. 2016 |
|  | *Fucus vesiculosus* | contraction south | Nicastro et al. 2013 |
|  |  | expansion southwards | Lima et al. 2007 |
| Tree | *Thuja occidentalis* | expansion northwards | Boisvert & Périé 2014 |
|  |  | expansion northwards | Fei et al. 2017 |
|  |  | contraction north | Sittaro et al. 2017 |
|  | *Pinus sylvestris* | expansion southwards | Kuhn et al. 2016 |
|  | *Juniperus communis* | expansion northwards | Groom 2013 |
|  | *Taxus baccata* | expansion northwards | Amano et al. 2014 |

The type of range margin (HLM or LLM) explained only a moderate 4% of the overall variation in the relative performance of marginal populations. This is unsurprising given the great variety of organisms, response variables, and ecological contexts considered in our analysis. In addition, most primary studies reported only short-term data that are likely to miss relevant periods of (by definition rare) climatic extreme events and may thus not fully capture long-term trends in those populations. Finally, performance at certain specific life stages is not necessarily a reliable predictor of lifetime fitness and population growth rates (Villellas et al. 2015, Lee-Yaw et al. 2016). Despite these diverse limitations, the type of range margin still was the main predictor of performance in marginal populations. Thirty-seven (82%) of the 45 comparisons available for the low-latitude margin showed worse performance in LLM than central populations, compared to 34 (53%) of 64 comparisons for HLM populations.

Our findings suggest that latitudinal asymmetries in demographic performance exist worldwide, and occur in both animals and plants and in both terrestrial and marine species. This ubiquity is particularly striking given the great diversity of ecological strategies to cope with environmental stresses and hazards. For instance, plants generally have a greater capacity to buffer climatic stress through phenotypic plasticity and persistent life cycle stages than animals (Villellas et al. 2015, see also Lloret et al. 2012), which would allow them to slow population declines and accumulate a higher extinction debt (Jackson & Sax 2009, Jump et al. 2009). Moreover, climate is shifting at different pace in marine and terrestrial environments, with median temperatures increasing more than three times faster on land than at sea (Burrows et al. 2011). Water temperature and related properties typically drive population dynamics of marine species, whereas many LLM populations of terrestrial species are primarily constrained by water balance (Hampe & Jump 2011; Vilà-Cabrera et al. 2019). This difference may also explain why marine ectothermic animals tend to more fully occupy the latitudinal ranges situated within their thermal tolerance limits than terrestrial ectotherms (Sunday et al. 2012).

Although purely correlational, the analysis of relationships between differences in population performance and local climates provides interesting insights that add further support to our climate-change based interpretation of geographical trends in marginal population performance. First, centre-margin differences in population performance were generally related to differences in temperature, yet this link was far stronger in LLM populations than in HLM populations. Such an asymmetry is expected under recent global climate warming which tends to exacerbate temperature-related climatic constraints for LLM population performance, while relaxing them in HLM populations (Normand et al. 2009, Hastings et al. 2020, Kuntsler et al. 2020). Low levels of precipitation reinforced the observed temperature effect, and this was once again especially true in LLM populations. This trend is likewise expectable under recent climate warming, given that many LLM populations of terrestrial organisms experience constraints from water availability (Vilà-Cabrera et al. 2019) and their performance should hence suffer most strongly when a temperature increase goes along with low levels of precipitation.

## Conclusions

Given that differences in population performance can represent a powerful early indicator of impending range shifts (Parmesan et al. 1999, Lenoir & Svenning 2015), our results indicate that many extant species ranges are not in equilibrium with current climates even though they to date have not experienced perceivable shifts. Considering empirical fitness trends in marginal populations will substantially increase the realism of population-based approaches to species distribution modelling (Anderson et al. 2009, Maier et al. 2014). Given that latitudinal range shifts are likely to be ongoing or impending for many species, such improved predictive capacity is needed if we are to forecast their implications for biodiversity and ecosystem function.

## Acknowledgments

We are grateful to Amy L. Angert and Sergei Volis for supplying unpublished information, and to Pedro Jordano for insightful comments on the manuscript. This study was funded by NordForsk grant no. 80167 to the NORA consortium (Nordic Network for the Study of Species Range Dynamics, 2009–2012), by projects POPULIM (CGL2010-22180) and PERSLIM (CGL2010-18381) of the Spanish MICINN, the EU BiodivERsA project BeFoFu (NE/G002118/1) and the INRA ACCAF project FORADAPT. FRS was funded by a postdoctoral fellowship from the Spanish Ministerio de Economía y Competitividad (FPD2013-16756) and a Talent Attraction grant from the VI Plan Propio de Investigación at Universidad de Sevilla (VIPPIT-2018-IV.2). JCS considers this work a contribution to his VILLUM Investigator project (VILLUM FONDEN, grant 16549) and his European Research Council project (ERC-2012-StG-310886-HISTFUNC).

## Data and materials availability

Data and code that support the findings of this study have been archived at https://doi.org/10.15454/STOJ93.

## Appendix 1.

List of the 51 papers contributing data on population performance for the analysis.

## Appendix 2.

References cited in Table 2.

## Notes

### Competing Interest Statement

The authors have declared no competing interest.

