## Supplementary Material for "Widespread latitudinal asymmetry in marginal population performance"

### S1. Bibliographic compilation

We searched the ISI Web of Science (WOS) and Scopus until 19th April 2020 for papers containing adequate data for our study. The terms 'centre/center AND periphery', 'central AND marginal', 'abundant centre/center' and 'latitudinal cline' were introduced in combination with performance related terms including 'fecundity', 'survival', 'recruitment' and 'population growth rate'. We restricted our search to the categories: Environmental Sciences-Ecology, Plant Sciences, Zoology, Entomology, Marine and Freshwater Biology, Biodiversity Conservation, Agriculture, and Forestry in WOS ('Theme' as the search field) and Agricultural and Biological Sciences and Environmental Science in Scopus ('Article title, abstract and keywords' as search fields). We performed an additional search with Google Scholar (which tends to generate a larger number of papers but lacks specific tools for search refinement) by combining the search terms with the term 'ecology' and restricting our screening to the first 200 papers found.

Then we screened their abstracts or, when necessary, the main text of the articles to select only those papers fulfilling our criteria: (1) we only considered studies reporting field data from natural populations (including control populations of transplant experiments if these were measured at their home sites and met all other criteria); (2) we only considered studies with at least two central and two peripheral populations (i.e., true replicates); and (3) we only considered papers that provided sufficiently clear criteria for the definition of central and peripheral range parts relative to the global range of the target species. Finally, we searched the text of the selected papers and came to a final set of 51 papers that provided data amenable to meta-analysis, either primary data or data extracted from figures. These papers were then classified in two major kinds. First, papers including explicit centre-periphery comparisons of mean performance values from populations described as central or peripheral in the text. Second, papers based on latitudinal clines. In this later case, from each region we used the three most central and the three most extreme populations along the gradient (or more when several populations were located closely together).

### S2. Meta-analysis

#### *S2.1. Is the dataset subject to publication bias?*

Publication bias occurs when a dataset lacks disproportionately many case studies with either positive or negative effect sizes, that is, when some tendency has been more likely to be published (publication bias) or retrieved (dissemination bias) than others. We used four complementary approaches to estimate whether publication bias was likely to occur in our dataset: (1) visual inspection of a funnel plot, (2) the calculation of a fail-safe number, (3) a correlation between reported effect sizes and the impact factor of source journals, and (4) a cumulative meta-analysis to test for time-lag bias.

*(1) Funnel plot.* Funnel plots probe whether studies with little precision (small sample size studies) give different results from studies with greater precision (larger

studies). Asymmetry in the funnel plot is often interpreted as a sign of publication bias (i.e., the decision of authors or editors to publish or not a given result) or dissemination bias (i.e., small studies tend to be published in poorly accessible or indexed journals). On the contrary, the funnel plot we constructed from our dataset was symmetrical, indicating that small and large studies, as well as studies reporting negative, positive or close to zero effect sizes were equally likely to be published.

The utility of funnel plots in the context of multi-level meta-analysis remains a matter of debate, because sets of points may be clustered together as a result of statistical dependencies. However, there was no evidence for such clustering, and data points corresponding to HLM and LLM case studies were fairly well distributed. This observation further supports our conclusion that publication bias was unlikely in our dataset.

*(2) Fail-safe number.* Fail-safe numbers (FSN) estimate how many studies with effect sizes averaging zero should be added to negate the significance of the grand mean effect size (or to reduce it to a specified minimal value). Among the various available metrics, Koricheva *et al.* (2013) recommended the use of Rosenberg's FSN (2005), a weighted metric that is tested against a normal distribution. Rosenberg's FSN was 257 ($P < 0.001$), indicating that “publication biases (if they exist) may be safely ignored”.

*(3) Correlation between effect sizes and the impact factors of the reporting journal.* Publication bias is likely to occur if higher impact journals tend to publish papers with stronger effect sizes, whereas results reporting weaker or no empirical support for hypotheses are more likely published in lower rank (and maybe less accessible) journals (or not published at all). Following this logic, Murtaugh (2002) proposed a test of publication bias that consists of regressing effect sizes against the impact factor (IF) of the journal they were taken from. We used 2019 IF of journals that provided case studies included in the meta-analysis and assumed the rank of journal impact factors was stable over the period covered by our data. We found no correlation between effect sizes and IF ($r = -0.03$, $P = 0.744$). Strong effect sizes were neither more likely to be reported in top-rank journals, nor small effect sizes were more common in lower rank journals. Deleting case studies published in the journal Nature (IF > 40) did not change the conclusion ($r = -0.12$, $P = 0.249$).

*(4) Cumulative meta-analysis.* Temporal trends in effect sizes may affect the generality (and stability) of conclusions drawn from meta-analyses. Temporal trends may result from changes in methodology, technology or dominant paradigms. We assessed the temporal stability of the grand mean effect size (both for the complete dataset and for HLM and LLM separately) by conducting a cumulative meta-analysis. This analysis calculates the grand mean effect size of a successively accumulating subsample of the global dataset to which case studies are sequentially aggregated in their order of publication (i.e., from the oldest to the most recent publication year). We tested for the existence of a temporal trend by means of a weighted regression analysis with the year of publication as predictor variable and the grand mean effect sizes as response variable (function *rma.mv* in *metafor*).

Overall, grand mean effect sizes were fairly stable through time (Fig. S1). On the other hand, we observed a markedly stronger increase in the number of studies on HLM populations than on LLM populations through the past 10 years. Regardless of this difference, however, the difference between HLM and LLM remained strong and consistent, implying that our main result is likely to be insensitive to time-lag bias.

*S2.2. Accounting for potential non-independence of case studies drawn from the same primary paper*

Some primary papers contained more than one measure that we could use for our meta-analysis (e.g., reporting different performance estimators for the same species or the same estimator for different species). Such measures could be mutually non-independent, leading to pseudo-replication in the dataset used for the meta-analysis.

We used two complementary approaches to account for potential non-independence of case studies stemming from the same primary paper. (1) We used multi-level meta-analysis where we specified two random factors with the rma.mv function of the R package *metafor* (Viechtbauer 2010): the identity of the case study and the identity of the primary paper; the first factor was nested within the second. (2) We ran a sensitivity analysis to assess the robustness of the main result (i.e., margins differ in relative population performance) against the non-independence of case studies from the same primary paper.

*Sensitivity analysis*: we created a subsample of our global dataset that contained only one randomly selected case study from each primary paper. We then performed a mixed-effect meta-analysis on this subsample to test for the existence of a margin (HLM vs. LLM) effect with the rma function from package *metafor* (Viechtbauer 2010). This procedure was repeated 1000 times, each time with a newly created test dataset (which is analogous to bootstrap procedures using random drawing with replacement). Among the 1000 models, 35.4% supported a significant difference between HLM and LLM. The mean (± 95% distribution) of random samples for HLM (-0.06 ± [-0.45, 0.31]) and LLM (-0.87 ± [-1.37, -0.36]) were very close to model parameters estimated from the multi-level error meta-analyis presented in the main text (-0.37 ± [-0.80, 0.07] and -0.94 ± [-1.42, -0.46]) for HLM and LLM, respectively; see also Fig. 2). The sensitivity analysis thus supports our main finding, implying that the reported asymmetry in the relative performance of HLM and LLM populations is unaffected by potential lack of independence among case studies stemming from the same primary paper.

*S2.3. Does asymmetry in marginal population performance differ between taxonomic kingdoms (animals vs. plants) and between major habitats (terrestrial vs. marine)?*

Our model selection procedure retained five models within two units of ΔAICc of the best model. All included margin type as a moderator and the null model (i.e., intercept only) was excluded (Table S1, S2**.**

To assess the relative relevance of each of our three moderators, we calculated the sum of weights ($w_H$) of individual moderators as the sum of weights of all models ($w_i$) with this predictor and ΔAICc < 10 (Burnham & Anderson 2004). The result is shown in Fig. S2.

Margin type was the most important predictor ($w_{H,\ margin}$ = 0.99), whereas Habitat ($w_{H,habitat}$ = 0.73) and Kingdom ($w_{H,\ kingdom}$ = 0.48) received only marginal support, and interactions (Margin × Habitat and Margin × Kingdom) were even less relevant. Fully in line with this result, neither Kingdom nor Habitat explained a significant amount of heterogeneity in any of the five models retained in the set of best models (Table S2).

The combined evidence supports our conclusion that the reported latitudinal asymmetry in marginal population performance is not discernibly affected by differences in effect sizes between plants and animals or between marine and terrestrial organisms.

### S3. Description and results of the climate analysis

The relationship between the relative performance at marginal populations - (Hedges' *d*) and the difference in average climate between marginal and central populations in the period 1985–2016 was analyzed by means of generalized additive mixed models (GAMM). In particular, we used the following model:

*RelativePerformance ~ s(TemperatureDifference) + s(study, bs* = “re”)

where *s* represents smooth terms (Wood 2006), in this case using thin plate splines. We used random effects smooths (*bs* = "re") to account for non-independence of comparisons within published studies. We also weighted relative performance effect sizes (Hedges' *d*) by their variances so that their influence in model calibration was inversely related to their uncertainty (Rayner et al. 2003).

We found a moderate but statistically significant effect of temperature on the relative performance of marginal populations (estimated degrees of freedom = 3.08, $P$ = 0.017, Table S3). The model managed to explain 21% of the total deviance.

For terrestrial species, we also fitted a similar model including precipitation and its interaction with temperature as predictors:

*RelativePerformance ~ s(TemperatureDifference) + s(PrecipitationDifference) + ti(TemperatureDifference, PrecipitationDifference) + s(study, bs* = “re”)

where *PrecipitationDifference* is the difference in average annual precipitation between marginal and central populations, and *ti* accounts for the tensor product interaction between temperature and precipitation.

In terrestrial species, population performance showed a similar asymmetric response to temperature (i.e., higher overall performance in HLM than LLM for similar temperature deviations), although there was a significant interactive effect of precipitation ($F$ = 2.695, $P$ = 0.024, total deviance explained = 29.5%; Table S4).

We fitted these models using package mgcv v. 1.8-33 (Wood 2006) in R 4.0.3 (R Core Team 2016). The entire code to reproduce these analyses is available as a research compendium in Rodríguez-Sánchez (2017).

### Supplementary tables

**Table S1.** Summary of models with ΔAICc, $w_i$ and heterogeneity. Model selection procedure retained five models within two units of ΔAICc of the best model (bold characters). All included margin type as a moderator and the null model (i.e., intercept only) was excluded. Only models with ΔAICc < 10 are presented. Pseudo $R^2$ was calculated as 1 – LLR where LLR is the ratio between the log-likelihood of model $i$ and the log-likelihood of the null model.

| Model | AICc | ΔAICc | wi | QM | P(QM) | QE | P(QE) | Pseudo R² |
|---|---|---|---|---|---|---|---|---|
| HABITAT + MARGIN | 455.75 | 0.00 | 0.20 | 14.41 | 0.001 | 335.62 | < 0.001 | 0.040 |
| MARGIN | 456.05 | 0.30 | 0.18 | 11.74 | 0.001 | 342.87 | < 0.001 | 0.029 |
| HABITAT + MARGIN + MARGIN:HABITAT | 456.53 | 0.78 | 0.14 | 15.75 | 0.001 | 334.47 | < 0.001 | 0.048 |
| HABITAT + MARGIN + KINGDOM | 456.82 | 1.07 | 0.12 | 15.73 | 0.001 | 334.39 | < 0.001 | 0.048 |
| HABITAT + MARGIN + KINGDOM + MARGIN:HABITAT | 457.39 | 1.63 | 0.09 | 17.26 | 0.002 | 332.96 | < 0.001 | 0.057 |
| MARGIN + KINGDOM | 458.22 | 2.47 | 0.06 | 11.75 | 0.003 | 342.58 | < 0.001 | 0.035 |
| HABITAT + MARGIN + KINGDOM + KINGDOM:MARGIN | 458.54 | 2.78 | 0.05 | 16.34 | 0.003 | 329.90 | < 0.001 | 0.054 |
| HABITAT + MARGIN + KINGDOM + KINGDOM:HABITAT | 458.99 | 3.24 | 0.04 | 15.85 | 0.003 | 334.18 | < 0.001 | 0.054 |
| HABITAT + MARGIN + KINGDOM + MARGIN:HABITAT + KINGDOM:HABITAT | 459.38 | 3.63 | 0.03 | 17.61 | 0.003 | 332.39 | < 0.001 | 0.063 |
| MARGIN + KINGDOM + KINGDOM:MARGIN | 459.52 | 3.77 | 0.03 | 12.90 | 0.005 | 337.14 | < 0.001 | 0.041 |
| HABITAT + MARGIN + KINGDOM + MARGIN:HABITAT + KINGDOM:MARGIN | 459.69 | 3.94 | 0.03 | 17.29 | 0.004 | 329.90 | < 0.001 | 0.061 |
| HABITAT + MARGIN + KINGDOM + KINGDOM:HABITAT + KINGDOM:MARGIN | 460.78 | 5.03 | 0.02 | 16.42 | 0.006 | 329.84 | < 0.001 | 0.060 |

| | | | | | | | | |
|---|---|---|---|---|---|---|---|---|
| HABITAT + MARGIN + KINGDOM + MARGIN:HABITAT + KINGDOM:HABITAT + KINGDOM:MARGIN | 461.74 | 5.99 | 0.01 | 17.61 | 0.007 | 329.82 | < 0.001 | 0.068 |
| Null | 465.26 | 9.51 | 0.00 | - | - | 362.48 | < 0.001 | 0.000 |
| HABITAT | 465.30 | 9.55 | 0.00 | 2.27 | 0.132 | 356.65 | < 0.001 | 0.010 |
| HABITAT + KINGDOM | 466.68 | 10.92 | 0.00 | 3.18 | 0.203 | 356.11 | < 0.001 | 0.018 |
| KINGDOM | 467.41 | 11.66 | 0.00 | 0.00 | 0.972 | 362.00 | < 0.001 | 0.006 |
| HABITAT + KINGDOM + KINGDOM:HABITAT | 468.90 | 13.15 | 0.00 | 3.19 | 0.364 | 356.01 | < 0.001 | 0.024 |

**Table S2. Summary of the five models retained in the set of best models** (i.e., with ΔAICc < 2). Margin explained a significant amount of heterogeneity in each of the five competing best models whereas neither Kingdom nor Habitat explained a significant amount of heterogeneity in any of the five models retained in the set of best models. For each moderator, coefficient parameter estimates are compared to the intercept. $Q_M$ and associated *P-values* represent the omnibus test. Pseudo $R^2$ were calculated as 1 – LLR, where LLR is the ratio between the log-likelihood of model *i* and the log-likelihood of the null model.

| **Model** | **Parameters** | **Estimate (± 95% CI)** | **Z-value** | **P-value** | **QM (P-value)** | **QE (P-value)** | **Pseudo $R^2$** |
|---|---|---|---|---|---|---|---|
| Habitat + Margin | Intercept ($Margin_{HLM}$, $Habitat_{Marine}$) | 0.3 (-0.28, 0.88) | 1.02 | 0.309 | 14.06 (0.001) | 335.62 (< 0.001) | 0.04 |
| | $Margin_{LLM}$ | -0.72 (-1.12, -0.31) | -3.47 | < 0.001 | | | |
| | $Habitat_{Terrestrial}$ | -0.5 (-1.14, 0.14) | -1.55 | 0.122 | | | |
| Margin | Intercept ($Margin_{HLM}$) | -0.07 (-0.41, 0.27) | -0.41 | 0.681 | 11.65 (0.001) | 342.87 (< 0.001) | 0.029 |
| | $Margin_{LLM}$ | -0.71 (-1.11, -0.3) | -3.41 | < 0.001 | | | |
| Habitat + Margin + Habitat × | Intercept ($Margin_{HLM}$, $Habitat_{Marine}$) | 0.49 (-0.17, 1.15) | 1.45 | 0.148 | 15.45 (0.001) | 334.47 (< 0.001) | 0.048 |

| | | | | | | | |
|---|---|---|---|---|---|---|---|
| Margin | | | | | | | |
| | Margin$_{\text{LLM}}$ | -1.16 (-1.95, -0.36) | -2.85 | 0.004 | | | |
| | Habitat$_{\text{Terrestrial}}$ | -0.75 (-1.51, 0.01) | -1.94 | 0.053 | | | |
| | Margin$_{\text{LLM}}$ × Habitat$_{\text{Terrestrial}}$ | 0.59 (-0.33, 1.5) | 1.26 | 0.209 | | | |
| Habitat + Margin + Kingdom | Intercept (Margin$_{\text{HLM}}$, Habitat$_{\text{Marine}}$, Kingdom$_{\text{Animals}}$) | 0.23 (-0.36, 0.82) | 0.77 | 0.441 | 15.07 (0.002) | 334.39 (< 0.001) | 0.049 |
| | Margin$_{\text{LLM}}$ | -0.73 (-1.13, -0.32) | -3.51 | < 0.001 | | | |
| | Habitat$_{\text{Terrestrial}}$ | -0.69 (-1.43, 0.04) | -1.86 | 0.063 | | | |
| | Kingdom$_{\text{Plants}}$ | 0.35 (-0.32, 1.02) | 1.02 | 0.306 | | | |
| Habitat + Margin + Kingdom + Habitat × Margin | Intercept (Margin$_{\text{HLM}}$, Habitat$_{\text{Marine}}$, Kingdom$_{\text{Animals}}$) | 0.43 (-0.24, 1.09) | 1.26 | 0.208 | 16.69 (0.002) | 332.96 (< 0.001) | 0.057 |
| | Margin$_{\text{LLM}}$ | -1.21 (-2.01, -0.41) | -2.95 | 0.003 | | | |
| | Habitat$_{\text{Terrestrial}}$ | -0.99 (-1.85, -0.13) | -2.26 | 0.024 | | | |
| | Kingdom$_{\text{Plants}}$ | 0.4 (-0.29, 1.08) | 1.14 | 0.254 | | | |
| | Margin$_{\text{LLM}}$ × Habitat$_{\text{Terrestrial}}$ | 0.64 (-0.28, 1.56) | 1.37 | 0.172 | | | |

**Table S3. Results of the additive mixed model** relating relative performance of marginal populations (Hedges' *d*) to the difference in average climate between marginal and central populations. We used temperature difference as predictor, and the study as random effect (see Supplementary Information S3).

| | **Estimates** |
|---|---|
| (Intercept) | -0.31 (0.14)* |
| EDF: s(Tmean.dif) | 3.08 (3.80)* |
| EDF: s(study.id) | 4.28 (48.00) |
| Deviance explained | 20.8% |
| $R^2$ | 0.151 |
| Num. obs. | 109 |

*$p < 0.05$

**Table S4. Results of the additive mixed model** relating relative performance of marginal populations (Hedges' *d*) in terrestrial species to the difference in average climate between marginal and central populations. We used temperature and precipitation differences (and their interaction) as predictors, and the study as random effect (see Supplementary Information S3).

| | **Estimates** |
|---|---|
| (Intercept) | -0.36 (0.15)* |
| EDF: s(Tmean.dif) | 3.11 (3.85)* |
| EDF: s(Pmean.dif) | 1.00 (1.00) |
| EDF: ti(Tmean.dif, Pmean.dif) | 3.76 (4.79) |
| EDF: s(study.id) | 0 (34.00) |
| Deviance explained | 29.5% |
| $R^2$ | 0.219 |
| Num. obs. | 80 |

*$p < 0.05$

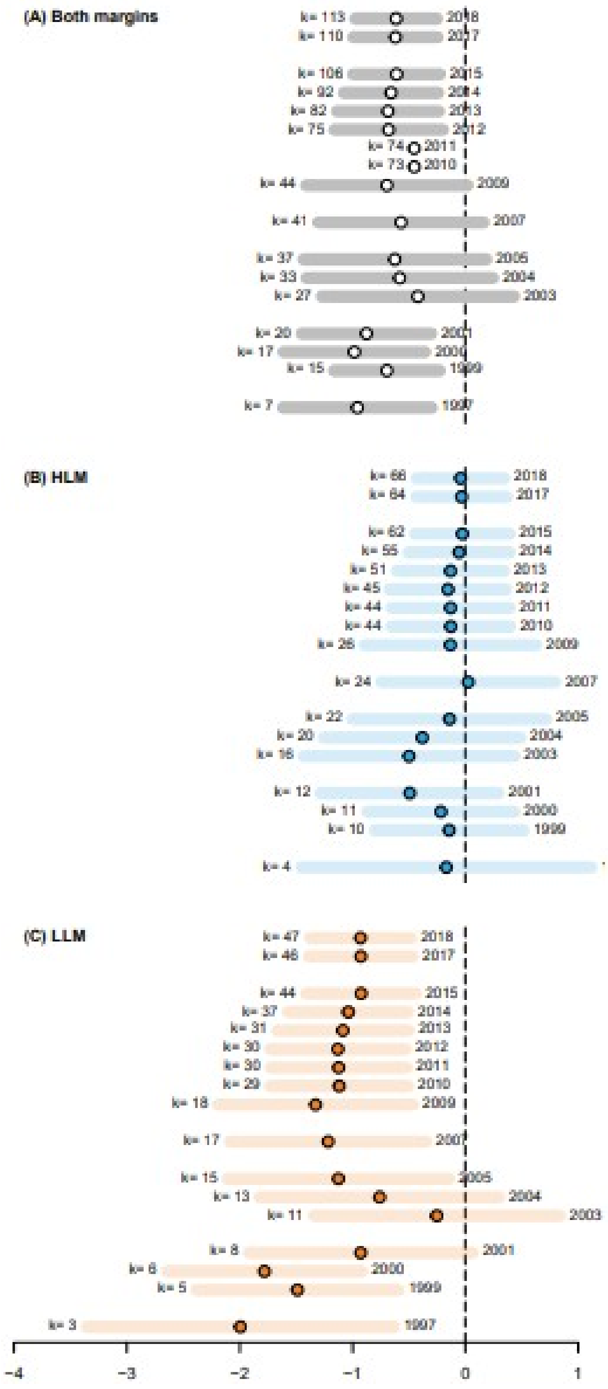


**Figure S1**. Cumulative meta-analysis. Grand mean effect sizes (dots), 95% CI (bars) and sample sizes (*k*) are shown for each year, including all previous years. Plate (A) depicts the global data set, plates (B) and (C) the datasets for HLM and LLM populations, respectively. Only significant relationships between publication year and effect sizes are shown by a regression line (continuous) and its 95% CI (dotted).

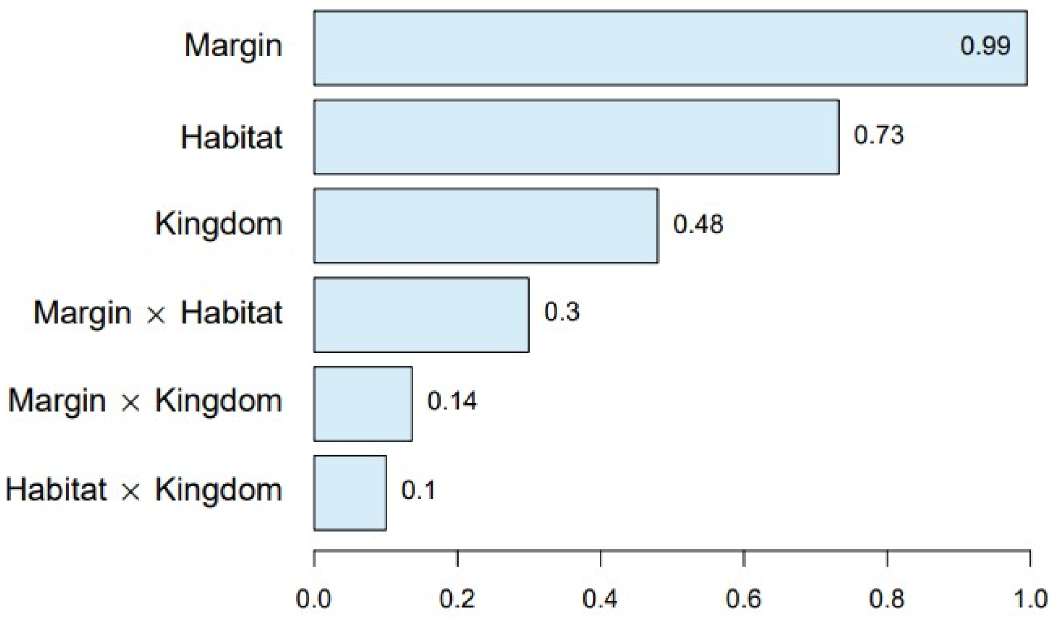


**Figure S2**. Sum of weights of moderators quantifying the relative importance of individual moderators and their interactions. Values are interpreted as the probability that a given variable is retained in the best model. Among tested moderators, margin type was the most important predictor (wH, margin = 0.96), whereas Habitat (wH,habitat = 0.69) and Kingdom (wH, kingdom = 0.54) received only marginal support, and interactions (Margin × Habitat and Margin × Kingdom) were even less relevant. Fully in line with this result, neither Kingdom nor Habitat explained a significant amount of heterogeneity in any of the five models retained in the set of best models (Table S2).

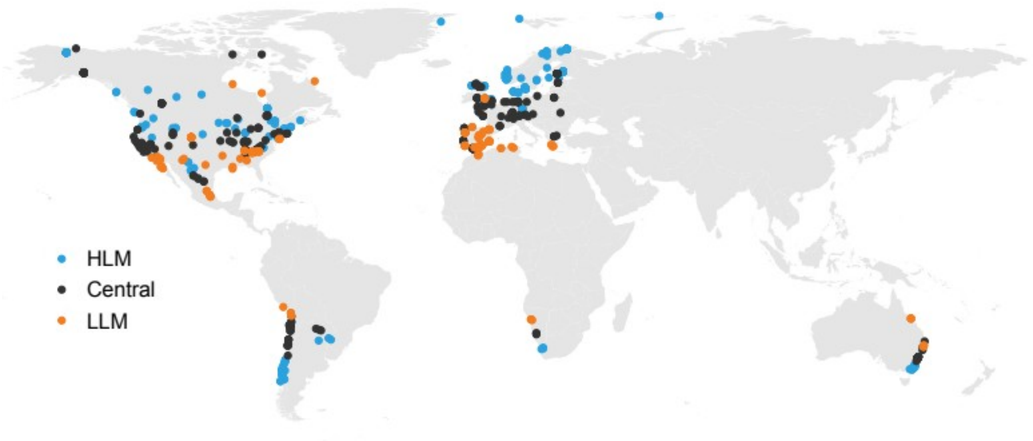


**Figure S3.** Map of the 687 populations included in this study, classified as 'High-Latitude Margin' (HLM), 'Central' populations, or 'Low-Latitude Margin' (LLM). Note that HLM populations for some organisms can be at lower latitudes than LLM populations of other species (and *vice versa*).

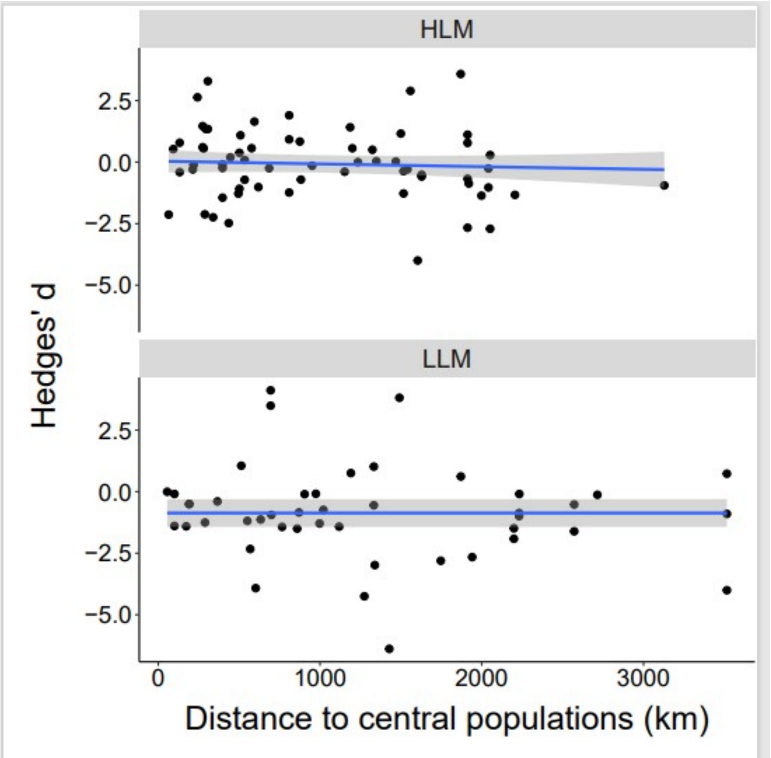


**Figure S4.** Relative performance of marginal vs central populations (Hedges' *d*) in relation to the geographic distance between them. The latter was calculated as the distance between the centroids of marginal (HLM or LLM) and central populations in each case. We found no evidence for a distance effect on explaining differences in relative population performance, as we found for climate (Fig. 4).
